# Loss of *dop-2* causes increased dopamine release and locomotory defects in the presence of ethanol

**DOI:** 10.1101/779405

**Authors:** Pratima Pandey, Anuradha Singh, Harjot Kaur, Anindya Ghosh-Roy, Kavita Babu

## Abstract

Ethanol is a widely used drug, excessive consumption of which could lead to medical conditions with diverse symptoms. Ethanol abuse causes disinhibition of memory, attention, speech and locomotion across species. Dopamine signaling plays an essential role in ethanol dependent behaviors in animals ranging from *C. elegans* to humans. We devised an ethanol dependent assay in which mutants in the dopamine autoreceptor, *dop-2,* displayed a unique sedative locomotory behavior causing the animals to move in circles while dragging the posterior half of their body. We identify the posterior dopaminergic sensory neuron as being essential to modulate this behavior. We further demonstrate that in *dop-2* mutants, ethanol exposure increases dopamine secretion and results in enhanced function of the DVA interneuron. DVA releases the neuropeptide NLP-12 and leads to the excitation of cholinergic motor neurons that affect movement. Thus, DOP-2 modulates dopamine levels at the synapse and regulates alcohol induced movement through NLP-12.

## Introduction

Alcohol is an easily available abusive drug used world over. Since excessive alcohol intake is detrimental to human health, many studies have focused on understanding the mode of action and dependency of this drug. Behavioral responses to alcohol and susceptibility to alcohol use disorders (AUDs) vary since they are dependent upon environmental, physiological and genetic differences amongst individuals (Prescott and Kendler, 1999; Schuckit and Smith, 1996). Hence, it is still unclear how alcohol functions to modulate various behaviors, making it important to identify and analyze target gene/s and molecular pathways which functions to modulate behavioral phenotype/s upon alcohol intake.

AUDs require functioning through multiple synaptic molecules including acetylcholine (ACh), GABA, glutamate, dopamine (DA), neuropeptide-Y related pathways and ligand gated channels (Reviewed in (Bettinger and Davies, 2014; Harris and Trudell *et al*., 2008; Spanagel, 2009)). Ethanol (EtOH) intake has been shown to increase DA release which in turn induces the reward pathway and brings about disinhibition of behaviors (reviewed in (Baik, 2013)). The DA pathway comprises of two receptor subfamilies; D1-like (inhibitory) and D2-like (excitatory) receptors that function through the G-protein signaling pathway (Bunzow et al., 1988; Pandey and Harbinder, 2012; Zhou et al., 1990). A special class of D2 autoreceptors have also been identified and are located on dopaminergic neurons, these receptors are thought to regulate the release of DA (reviewed in (Ford, 2014)). In mammals, activation of these receptors lowers the excitability of DA neurons and modulates the release and transmission of dopamine (Beaulieu and Gainetdinov, 2011; Koeltzow et al., 1998; Rivet et al., 1994). Further, the D2 autoreceptors have been associated with alcoholism (Lu et al., 2001; Thanos et al., 2005).

*Caenorhabditis elegans* as a model organism is popular for its powerful genetic tools (Corsi et al., 2015) and has been widely utilized for studying the various aspects of neuroactive drugs (Alaimo et al., 2012; Bettinger et al., 2004; Giacomotto and Segalat, 2010; Hawkins et al., 2015; Schafer, 2004). *C. elegans* encounters alcohol in its natural habitat (rotten fruits) and hence could be expected to have evolutionarily developed neuronal circuitry allowing for alcohol sensitivity. The DA system is very compact in *C. elegans*, with merely eight DA neurons as compared to ∼500,000 neurons in the ventral midbrain of humans ((Sulston et al., 1975) and reviewed in (Hegarty et al., 2013)). *C. elegans* DA receptors, like their mammalian counterparts also belong to two subfamilies, D1-like and D2-like receptors (Suo et al., 2002, 2003). EtOH shows its effect in a concentration dependent manner, acting as stimulant at lower concentration and depressant at higher concentration. Studies in *C. elegans* have reported that EtOH administration shows dose dependent decline in the locomotor activity with increasing levels of EtOH exposure, which is similar to the depressive effects of EtOH seen in other animal systems (Alaimo et al., 2012; Davies et al., 2003; Hawkins et al., 2015). The internal dose of EtOH responsible for this behavior is similar to that in mammalian systems, indicating that there might exist similar molecular targets (Lee et al., 2009).

Utilizing *C. elegans*, we devised an EtOH dependent assay and screened for dopaminergic receptors and pathway mutants. Mutants in the D2-like autoreceptor, *dop-2* displayed a novel locomotory phenotype when exposed to 400 mM EtOH. DOP-2 has previously been shown to participate in associative learning and copulation behaviors (Correa et al., 2015; Voglis and Tavernarakis, 2008). We found that *dop-2* mutant animals slowly dragged their body in concentric circles in what we refer to as Ethanol Induced Sedative (EIS) behavior. Our experiments indicate that *dop-2* mutants show increased dopamine release in the presence of EtOH, which is responsible for the EIS phenotype seen in these animals. Previous work has implicated a circuit through the PDE sensory neurons, the DVA interneuron and cholinergic motor neurons that allow for normal locomotion in *C. elegans* (Bhattacharya et al., 2014; Hu et al., 2011). Our work builds on this previous circuit and goes on to elucidate that the same circuitry is hyperactivated in *dop-2* mutants in the presence of EtOH and this in turn causes the EIS behavior.

## Results

### Ethanol exposure affects movement in *dop-2* mutants

In *C. elegans,* the DA pathway is widely known to modulate egg laying, defecation, basal slowing, habituation, and associative learning (McDonald et al., 2006; Sawin et al., 2000; Voglis and Tavernarakis, 2008). DA receptors, DOP-1 and DOP-3 function antagonistically to regulate signaling in acetylcholine motor neurons (Allen et al., 2011), while the DA receptor, DOP-2 is expressed presynaptically in all the dopaminergic neurons and has the potential to function as an autoreceptor. However, the *dop-2* mutants do not show any obvious defects that can explain its autoreceptor function. It is possible that under wild-type conditions loss of *dop-2* is compensated for by other regulatory mechanisms in the organism. Modulation of behavior and function through the dopaminergic system upon exposure to drugs of abuse such as ethanol has been previously established (Kameda et al., 2007; Siciliano et al., 2019). We speculated that exposure of the mutants of the dopaminergic pathway to Ethanol (EtOH) might allow us to screen for possible behavioral defects in these mutants. Previous work has shown that wild type (WT) *C. elegans* show flattening of the body-bends at 400 mM concentration of EtOH (Davies and McIntire, 2004; Davies et al., 2003). We observed a similar phenotype with WT animals, but after a period of 2 hours (h) these animals recovered and began to move in a manner that was similar to WT animals that had not been exposed to EtOH (Fig. 1a-d and Movie 1). We went forward to screen mutants in the dopaminergic pathway for defects in locomotion beyond 2 hours in the presence of 400 mM EtOH and found that mutants in the dopamine autoreceptor, *dop-2*, showed a unique behavior where the *C. elegans* kept moving in circles and compulsively dragged the posterior part of their body (Fig. S1 and Movie 2). During this movement we observed that there was slowing of movement as measured by number of body-bends and a flattening of the body-bends (amplitude of body-bends) that was very pronounced in the posterior half of the animal (Figs. 1e and f). We referred to this behavior as an Ethanol Induced Sedative (EIS) behavior. In order to test if this behavior was seen only in the presence of EtOH or observable in untreated animals as well, we quantified the anterior and posterior body-bends and amplitude of body-bends from untreated WT and *dop-2* mutants and saw no significant difference between both the strains (Fig. 1g and h). We next wanted to test if *dop-2* mutants affect all muscles or largely affect the locomotory body-wall muscle function. To address this question we treated control WT animals, *dop-2* mutants and mutants in the *slo-1* BK Potassium channel that is required for alcohol sensing (Davies et al., 2003) with EtOH for more than 2 h and counted the number of pharyngeal pumps per minute (min) in each strain. We found that the pharyngeal pumping frequency of *dop-2* mutants was indistinguishable from that of WT animals (Fig. S2). These results suggest that *dop-2* mutants have higher, prolonged locomotory sedative response to EtOH.

**Fig. 1:**
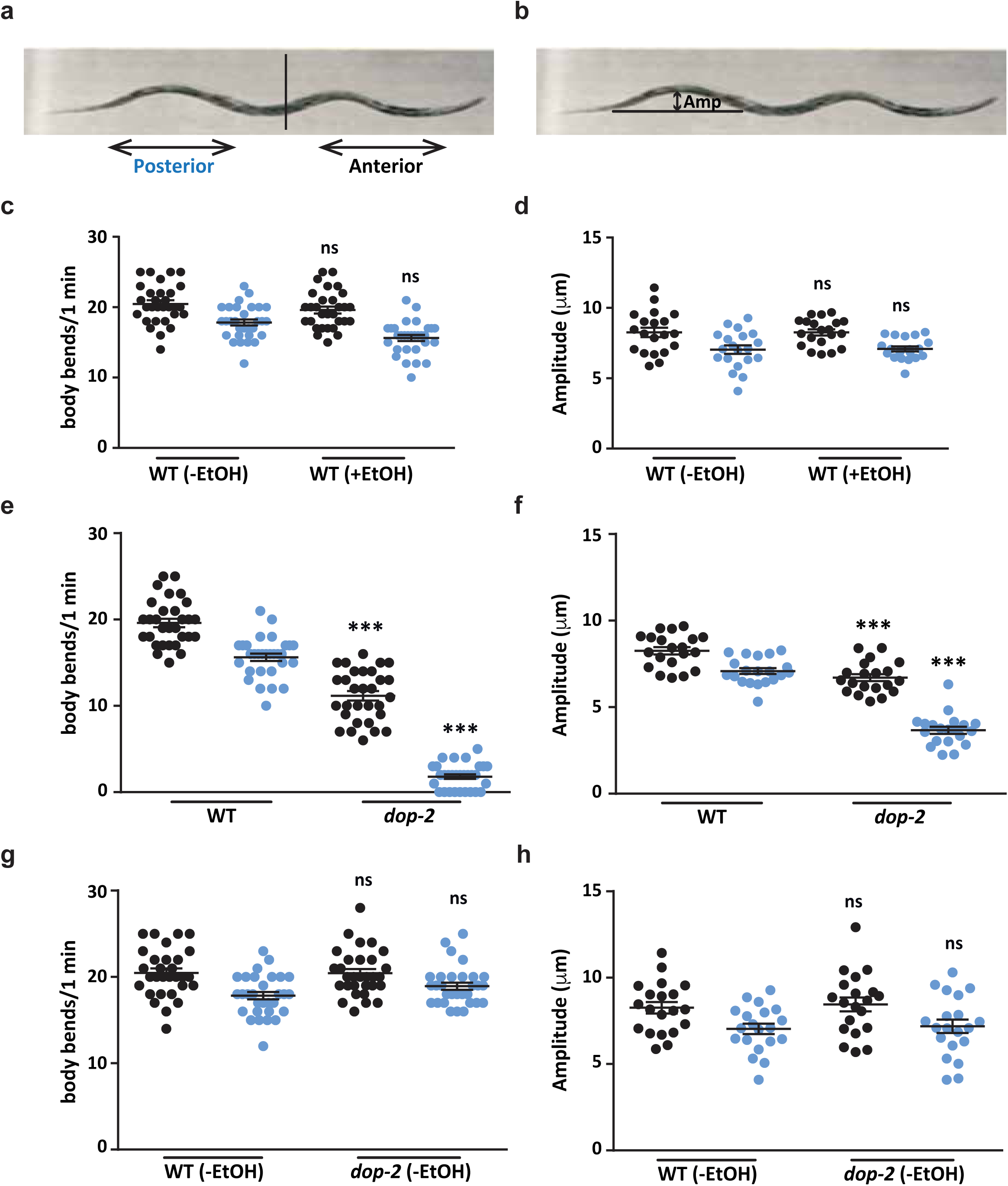
Analysis of movement defects upon exposure to 400 mM ethanol. (a) *C. elegans* body-bends in the anterior and posterior regions marked by a partition. (b) Amplitude of body-bends shown by a double-sided arrow. (c) Quantitative analysis of WT *C. elegans* for number of body-bends in case of untreated (-EtOH) and 400 mM Ethanol exposed animals (+EtOH). Body-bends were counted for 1 minute (min) after 2 hours (h) in both sets of animals (+/-EtOH). The number of body-bends per minute in the anterior and posterior region of the animal were plotted with posterior body-bends shown in blue in all figures. Experiments were carried out in triplicates with 10 worms each. In this graph F = 20.9 and DF = 3. (d) Quantitative analysis of amplitude of body-bends for WT untreated (-EtOH) and WT worms exposed to 400 mM EtOH (+EtOH). The amplitude of body-bends was analyzed for the animals using ImageJ and the measure is represented in μm. The amplitude of body-bends in the anterior and posterior regions of the animal were plotted with posterior body-bends shown in blue in all figures. The experiments were performed in duplicates and 20 worms were quantitated and represented in graphs. The same animals were used to quantitate body-bends and amplitude of body-bends for all figures. In this graph F = 6.97 and DF = 3. (e) Quantitative analysis of number of body-bends for WT and *dop-2* mutant worms, both treated with EtOH (F = 299, DF = 3). (f) Quantitative analysis of amplitude of body-bends for WT and *dop-2* mutant animals, both treated with EtOH (+EtOH) (F = 96.6, DF = 3) (g) Comparison of number of body-bends for WT and *dop-2* mutants in the absence of EtOH (-EtOH), (F = 7.57, DF = 3). (h) Quantitative analysis of amplitude of body-bends for WT and *dop-2* mutant worms in the absence of EtOH (-EtOH), (F = 4.09, DF = 3). Error bars represent ±S.E.M. and p-values were calculated using one-way ANOVA and Turkey-Kramer multiple comparison test; “***” indicates p<0.001 and “ns” indicates not significant in all graphs.

### The EIS behavior in *dop-2* is modulated through the PDE neuron

Our results suggest that EIS is a *dop-2* dependent behavior. To show that it is indeed dependent on DOP-2 we made transgenic rescue lines with the *Pdop-2*::DOP-2::CFP construct (Correa et al., 2012), and used this line in the EtOH assay and observed that both transgenic rescue lines could completely rescue the *dop-2* mutant phenotype (Figs. 2a and b). We next wanted to identify the neuron through which DOP-2 could be functioning.

**Fig. 2:**
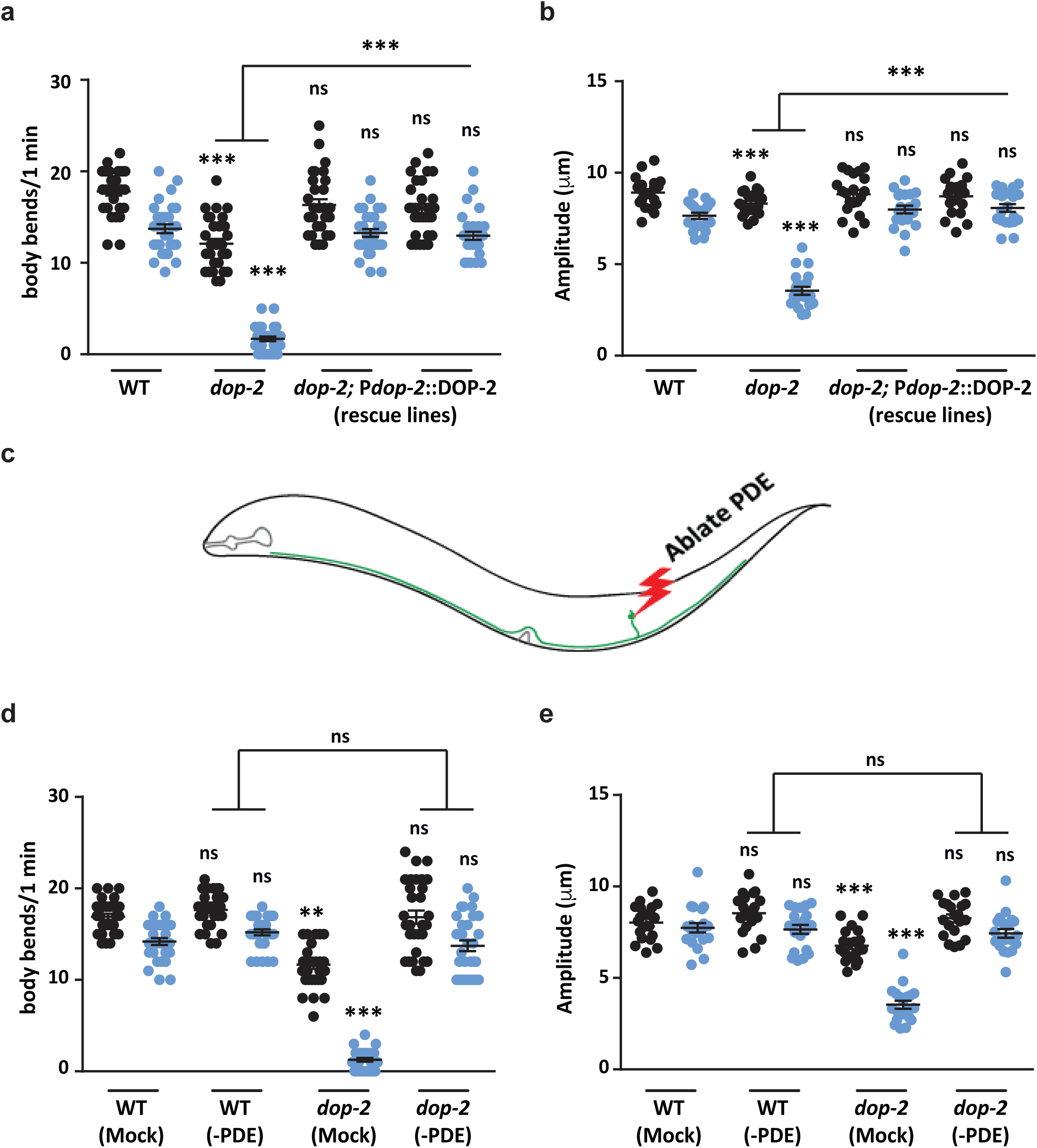
DOP-2 functions in the PDE neuron for the Ethanol Induced Sedetive (EIS) behavior. (a) Graph indicating body-bend measurements for rescue of the *dop-2* behavior using transgenic expression of DOP-2 under the *dop-2* promoter, (F = 109, DF = 7). (b) Graph indicates amplitude of body-bend measurements for WT, *dop-2* and rescue lines (F =74.6, DF = 7). (c) Illustration of PDE neuron ablation. (d) Quantitation of the number of body-bends in mock ablated and PDE ablated animals, (F =140, DF = 7) (e) Quantitation of the amplitude of body-bends in mock ablated and PDE ablated *C. elegans,* (F = 48.8, DF = 7). All experiments in this figure were performed in the presence of EtOH (+EtOH). Error bars represent ±S.E.M. and p-values were calculated using one-way ANOVA and Turkey-Kramer multiple comparison test; “**” indicates p<0.01, “***” indicates p<0.001 and “ns” indicates not significant in all panels.

There are eight dopaminergic neurons in the *C. elegans* hermaphrodite, two pair of CEP and a pair of ADE neurons in the head and a pair of PDE neurons in the posterior half of the *C. elegans* body (Sulston et al., 1975). All these dopaminergic neurons are mechanosensory in nature and control basal slowing behavior in the animal (Sawin et al., 2000). Our behavioral experiments indicated that the posterior half of the *dop-2* animals was more affected than the anterior in the presence of EtOH. The only dopaminergic neuron with sensory endings at the posterior region is the PDE neuron that is also involved in harsh touch behavior and context dependent modulation of movement (Bhattacharya et al., 2014; Li et al., 2011). Bhattacharya *et al* have recently shown that the PDE neuron through synaptic signaling via the DVA interneuron regulates the motor circuit in *C. elegans* (Bhattacharya and Francis, 2015; Bhattacharya et al., 2014). From the above information, we hypothesized that the EIS behavior could be due defective dopaminergic signaling from the PDE neuron. Since we were unable to find a PDE specific promoter for rescue experiments, we decided to ablate the PDE sensory neurons in WT and *dop-2* backgrounds and analyzed the PDE ablated animals for ethanol sensitivity (Illustrated in Fig. 2c). We observed that on exposure to EtOH the PDE ablated WT animals displayed no obvious defects in the number of body-bends or amplitude of body-bends when compared to the mock treated animals (Figs. 2d and e). However, upon testing the *dop-2* mutant animals in the EtOH assay we saw that unlike the mock ablated animals that showed the EIS behavior, the *dop-2* mutants with ablated PDE neurons behaved like control animals (Fig. 2d and e). These results indicate that the EIS behavior is DOP-2 dependent and that DOP-2 function in PDE neurons is sufficient to regulate this behavior.

### WT animals show EIS behavior in the presence of exogenous dopamine

Our data suggests that DOP-2 is functioning through the DA neuron, PDE and could be dependent on dopamine. D2 like autoreceptors have been shown to modulate the levels of dopamine through the regulation of transporters and components of the dopamine synthesis pathway (reviewed in (Ford, 2014)). However, the function of DOP-2 is still unclear. Since the EIS behavioral model provides us with an experimental system to investigate the function of DOP-2 in sedative movement during exposure to EtOH, hence we examined how DA levels and DA synthesis pathway components might affect the EIS behavior. To address this we utilized the *cat-2* mutants in the EtOH assay. CAT-2 encodes a tyrosine hydroxylase and is required to synthesize DA from tyrosine. The *cat-2 (n4547)* mutant used in this study is an allelic deletion and is reported to have 20-30% of WT levels of dopamine (Lints and Emmons, 1999; Sanyal et al., 2004). We performed the EtOH assay with this mutant of *cat-2.* Upon observation and quantitation of *cat-2* mutant behavior it was quite evident that decreased levels of dopamine are not involved in the sedative, sluggish behavior shown by *dop-2* (Figs. 1e, 3a and b). We next generated *cat-2; dop-2* double mutants and performed the EtOH assay with these animals. We observed that this double mutants showed a similar behavior as was seen in *cat-2* mutant animals (Figs. 3a and b). These results indicated that *cat-2* could be functioning upstream of *dop-2* and that the EIS phenotype was unlikely to be caused by decreased dopamine levels.

**Fig. 3:**
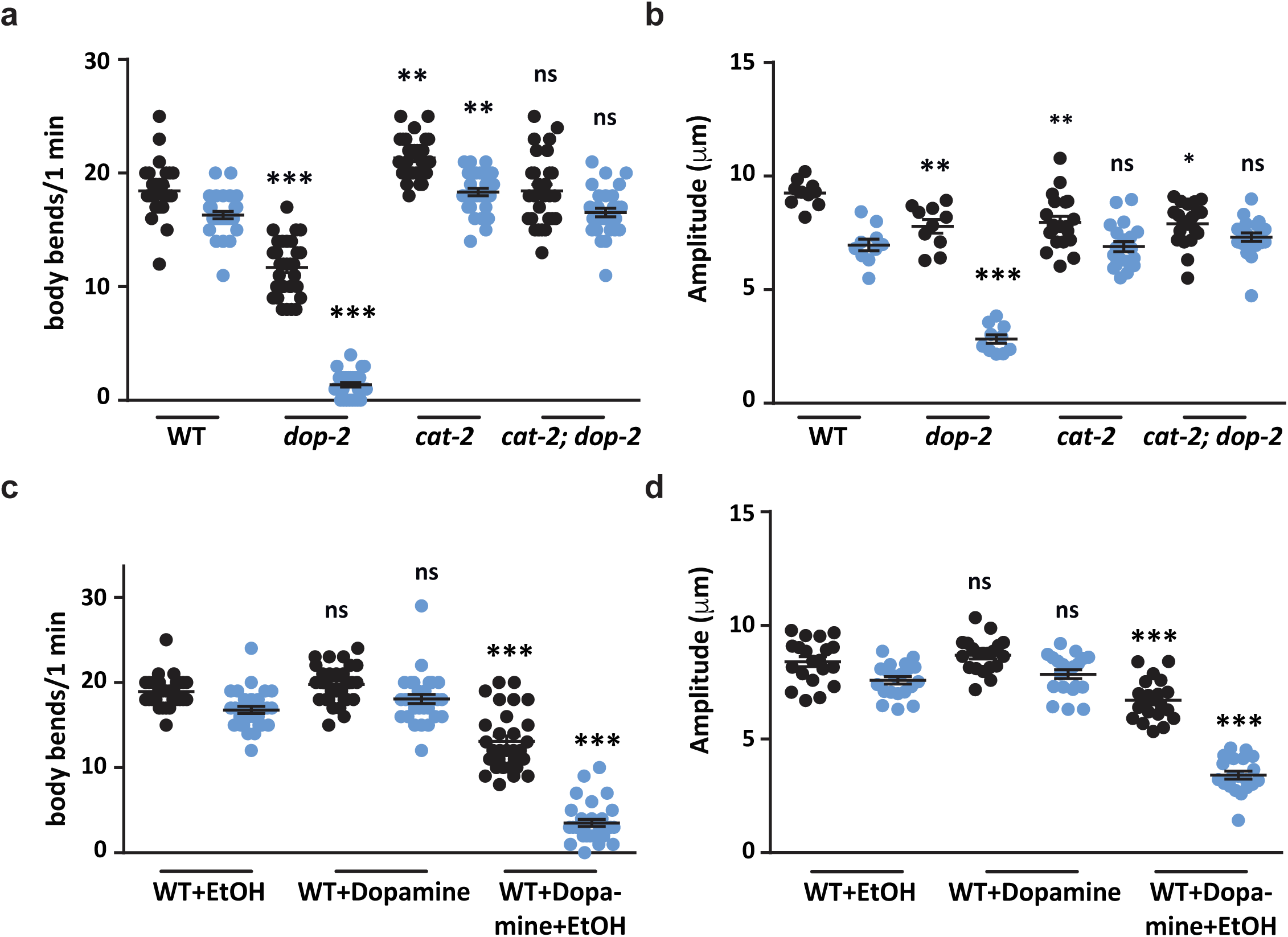
Role of dopamine in the ethanol induced behavior. (a) Graph represents the number of body-bends in WT, *dop-2, cat-2 and cat-2; dop-2* mutants in the presence of EtOH, (F = 265, DF = 7). (b) Quantitation of the amplitude of body-bends in WT, *dop-2*, *cat-2 and cat-2; dop-2* strains in the presence of EtOH, (F = 44.1, DF = 7). (c) Graph representing the number of body-bends in WT animals under different conditions (+ EtOH, + DA and + DA + EtOH), (F = 169, DF = 5). (d) Graph shows the amplitude of body-bends in WT *C. elegans* under different conditions (+ EtOH, + DA, + DA + EtOH), (F = 107, DF = 5). Error bars represent ±S.E.M and p-values were calculated using one-way ANOVA and Turkey-Kramer multiple comparison test; “*” indicates p<0.05, “**”indicates p<0.01, “***”indicates p<0.001 and “ns” indicates not significant in all panels.

There are multiple reports indicating the negative regulatory role of the D2 autoreceptor in regulating synaptic levels of dopamine (Benoit-Marand et al., 2001; Rouge-Pont et al., 2002; Schmitz et al., 2002). In order to test if the EIS phenotype was due to increased dopamine levels, we provided WT animals with exogenous dopamine in the EtOH assay. We observed that these animals showed the EIS behavior that was previously observed in *dop-2* mutants. These WT animals treated with exogenous dopamine showed decreased numbers and amplitude of body-bends in the EtOH plate. This phenotype was significantly different when compared to control WT animals treated with either just exogenous DA (no EtOH) or just EtOH (no exogenous DA) (Figs. 3c and d). These results suggest that increased dopamine release is responsible for the EIS behavior in *dop-2* mutant animals.

### Mutants in *dop-2* show increased dopamine release in the presence of EtOH

Our results so far have indicated that there could be increased levels of dopamine release in *dop-2* mutants in the presence of EtOH, which is responsible for the EIS behavior in these mutants. To get more insight into the defects in dopamine release in the PDE neurons of *dop-2* mutant animals treated with EtOH, we used FRAP (Fluorescence Recovery After Photobleaching) recordings as a tool to examine DA release (Miesenbock et al., 1998; Samuel et al., 2003). Here we used the previously constructed strain with synaptobrevin-super ecliptic pHluorin reporter fusion (SNB-1::SEpHluorin), which is expressed under the dopaminergic *asic-1* promoter ((Hardaway et al., 2015) and illustrated in Fig. S3). The PDE neuron synapses were examined with pH sensitive GFP (superecliptic pHluorin) attached to a vesicular protein SNB-1. The fluorescence was bleached at the synapses of PDE neuron and the rate of recovery at the bleached area was calculated as a possible measure of DA release. Increased release of dopamine was monitored by the rate of recovery in PDE synapses post bleach. We observed that EtOH exposed *dop-2* mutants showed a significantly faster rate of recovery in the case of *dop-2* mutant animals in presence of EtOH (Figs. 4a and b) and increased rates of fluorescence recovery at 60 s and 120 s time points in *dop-2* animals treated with EtOH (Fig. 4c). We next wanted to test if *dop-2* mutants affect levels of the surface DA transporter DAT-1.

**Fig. 4:**
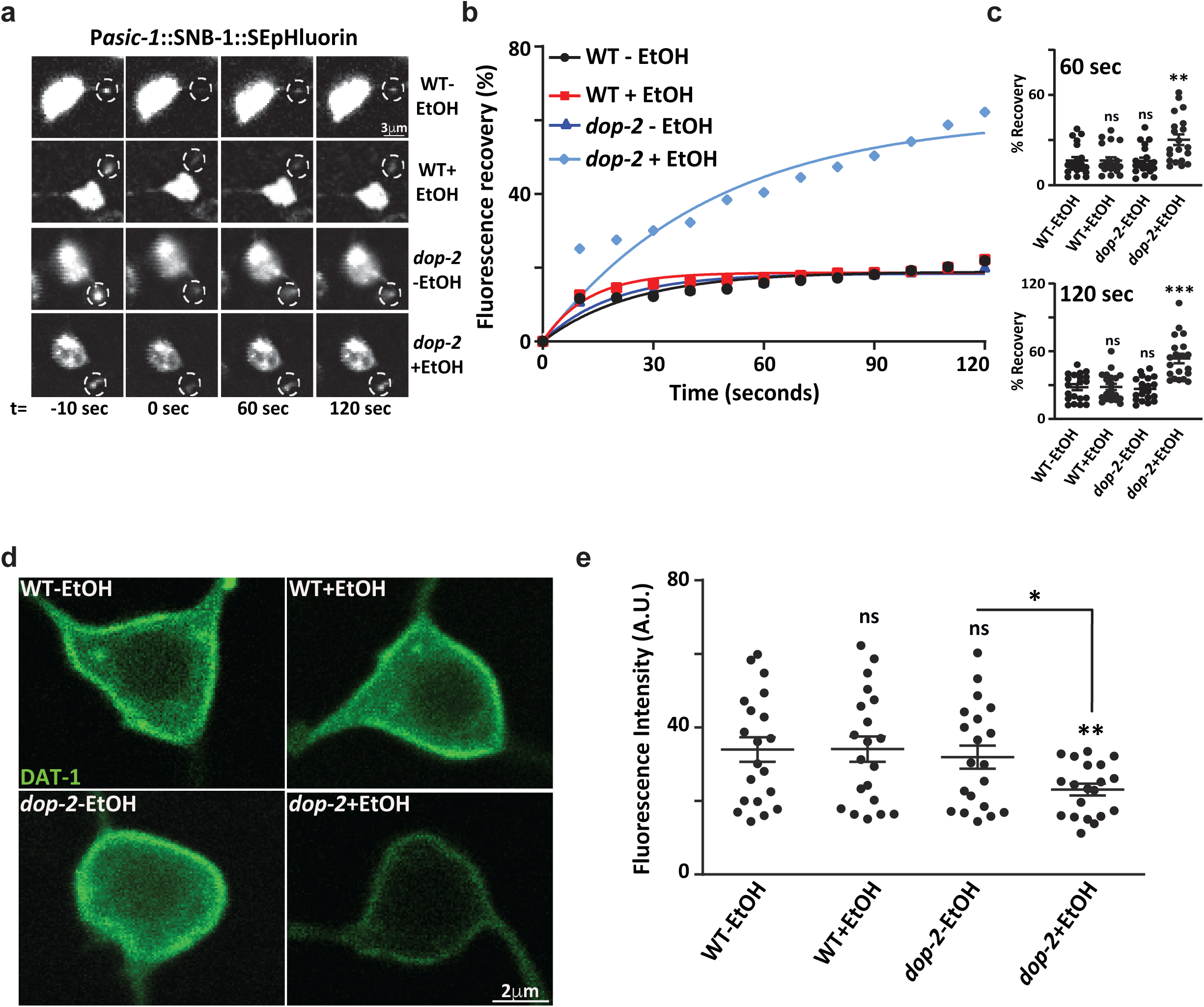
Mutants in *dop-2* show increased dopamine release in presence of ethanol. (a) Fluorescence recovery after photobleaching (FRAP) was performed on the PDE neuron synapses labeled with P*asic-1*::SNB-1::SEpHluorin. Representative images before bleaching (−10 seconds (sec)), followed by bleaching (0 sec) and post bleach at 60 sec and 120 sec. (b) Quantitation of rate of recovery in WT and *dop-2* mutant backgrounds with and without (+/- EtOH) treatment over 120 sec FRAP time course. Data represents 20-22 synapses per genotype. (c) Representative dot plots from FRAP data for percentage recovery at 60 sec and 120 sec time points respectively. (d) Representative images of DAT-1::GFP expression in WT and *dop-2* mutant background with and without EtOH treatment. (e) Whole cell fluorescence quantification of DAT-1::GFP in PDE neurons for WT and *dop-2* mutants with and without EtOH (n=20). Error bars represent ±S.E.M., n represents the number of animals analyzed and p-values were calculated using one-way ANOVA and Turkey-Kramer multiple comparison test; “*” indicates p<0.05, “**”indicates p<0.01, “***” indicates p<0.001 and “ns” indicates not significant in all panels.

Previous reports indicate that D2 like receptors are involved in the surface localization of DAT-1. We utilized the previously constructed DAT-1::GFP translational fusion line to study DAT-1 expression in *dop-2* mutants (Carvelli et al., 2004). We performed imaging and quantitated the cell surface expression of DAT-1 transporter in PDE neuron in the presence and absence of EtOH in the WT and *dop-2* mutant backgrounds. We observed a reduction in cell surface expression of DAT-1 in *dop-2* mutants exposed to EtOH (Figs. 4d and e). These experiments indicate that *dop-2* mutants in the presence of EtOH show increased synaptic DA and decreased surface DAT-1.

DAT-1 is present on the DA neurons and recycles DA back into the neuron. We reasoned that if the reuptake mechanism was affected then *dat-1* (DA transporter) deletion mutant should also show EIS like phenotype since DA levels should be higher than WT, but that wasn’t the case (Fig. S1). It is possible that loss of *dat-1* is only a part of the phenotype that allows for the EIS behavior in the animals that is seen in *dop-2* mutants treated with EtOH.

### DOP-2 functions through DOP-1 present in the DVA neuron

Thus far our data indicates that the EIS behavior seen in *dop-2* animals is modulated by the neurotransmitter DA. PDE has been found to be responsible for the DA effect but how this leads to defects in movement is still unknown. A previous study has elegantly shown that DA released from PDE neurons can activate DOP-1 present on the DVA neuron ((Bhattacharya et al., 2014) and reviewed in (Bhattacharya and Francis, 2015)). The DVA neuron upon activation results in the release of neuropeptide NLP-12 that results in the activation of downstream motor neurons (Hu et al., 2011). The sensory neuron PDE forms strong synaptic connections with the interneuron DVA (Sawin et al., 2000). To understand the circuit through which PDE functions for the EIS phenotype we studied how the deletion of the dopaminergic receptor, *dop-1* could affect the movement in *C. elegans* after EtOH treatment. We found no significant differences in body-bends on comparing *dop-1* mutants with WT animals (Figs. 5a and b). Next we made *dop-2; dop-1* double mutants and observed that it showed the same kind of behavior as *dop-1* mutants (Figs. 5a and b). Thus, *dop-1* deletion was able to suppress the EIS phenotype seen in *dop-2* mutants. If deletion of *dop-1* is obstructing DA signaling from PDE to DVA then we reasoned that expressing DOP-1 specifically in DVA would restore the EIS phenotype in the *dop-2; dop-1* double mutant animals. We indeed found that expressing DOP-1 specifically in DVA using the *nlp-12* promoter made the *dop-2; dop-1* animals revert to the *dop-2* like EIS behavior (Figs. 5a and b). These data indicate that EIS is dependent upon DA released from PDE and the DOP-1 receptor present on the DVA neuron.

**Fig. 5:**
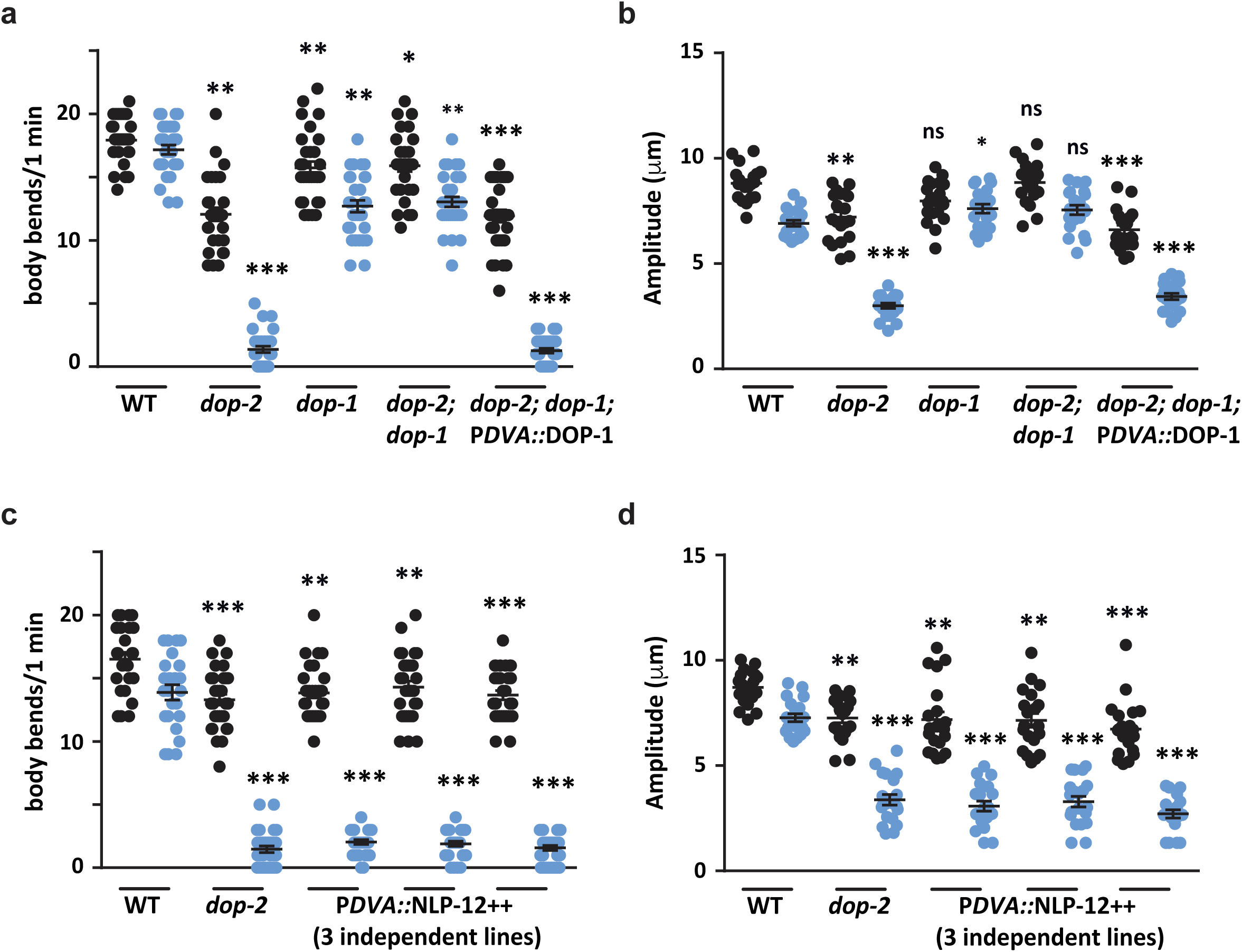
DOP-2 functions through the DOP-1 in the DVA neuron. (a) Graph shows number of body-bends in WT, *dop-2*, *dop-1*, *dop-2; dop-1* double mutants and the *dop-1* rescue line (*dop-2; dop-1*; P*DVA*::DOP-1) on EtOH treatment; (F = 212, DF = 9). (b) Graph shows amplitude of body-bends in WT, *dop-2*, *dop-1*, *dop-2; dop-1* double mutant and the *dop-1* rescue line (*dop-2; dop-1*; P*DVA*::DOP-1) on EtOH treatment, (F = 105, DF = 9). (c) Quantitation of number of body-bends in WT, *dop-2* and three NLP-12 overexpression lines upon EtOH treatment, (F = 302, DF = 9). (d) Quantitation of amplitude of body-bends in WT, *dop-2* and three NLP-12 overexpression lines upon EtOH treatment, (F = 77.2, DF = 9). Error bars represent ±S.E.M. and p-values were calculated using one-way ANOVA and Turkey-Kramer multiple comparison test; “*” indicates p<0.05, “**” indicates p<0.01, “***” indicates p<0.001 and “ns” indicates not significant in all panels.

The DVA neuron has been shown previously to function through the neuropeptide NLP-12. NLP-12 release potentially activates the downstream postsynaptic cholinergic motor neurons by binding to its receptors, CKR-2 (cholecystokinin like receptor). Interestingly, it has been shown previously that NLP-12 secretion is directly correlated with the speed of the animal (Hu et al., 2011). We reasoned that increased NLP-12 secretion could allow the animal to show the EIS phenotype. To perform this experiment we overexpressed NLP-12 in the DVA neuron. We found that all three NLP-12 overexpression lines showed the EIS behavioral phenotype (Figs. 5c and d). We also tested *nlp-12* mutants and found that they behaved in a manner similar to WT control animals after EtOH treatment (Figs. S4 a and b). We next wanted to test if ablating DVA would affect the EIS behavior in the NLP-12 overexpression line. However, we found that just ablating DVA in WT animals without EtOH treatment caused locomotory defects as has been shown previously ((Li et al., 2006), Figs. S4c and d). Our data so far suggests that dopamine released from PDE signals through DOP-1 receptors in the DVA interneuron, which in turn releases the neuropeptide NLP-12 in this circuitry is responsible for the EIS behavior in *dop-2* mutants treated with EtOH.

### Increased Acetylcholine levels at the NMJ results in EIS behavior

Previous work has shown that DA receptors DOP-1 (D1-like) and DOP-3 (D2-like) regulate locomotion in *C. elegans* (Chase et al., 2004; Omura et al., 2012). Studies also indicate that the hypercontracted state observed in case of ethanol exposure in the animals is due to increased acetylcholine at the NMJ (Hawkins et al., 2015). These studies prompted us to evaluate if dopamine might be involved in regulating locomotion through the cholinergic pathway. Initially we investigated if decreased levels of ACh could display the EIS behavior, but found no significant changes in the locomotion of cholinergic mutants when compared to WT control animals on exposure to EtOH (Fig. S5). We next wanted to test animals with increased ACh release in the presence of EtOH. Previous work has shown that aldicarb treatment causes increased ACh signaling at the NMJ (Hu et al., 2011). Hence, we performed the aldicarb assay followed by treatment of the animals with EtOH. For this experiment, *C. elegans* were exposed to 100 mM aldicarb followed by 400 mM EtOH exposure. We found that WT animals subjected to the above assay showed the EIS phenotype previously seen in *dop-2* mutant animals (Figs. 6a and b). The acetylcholine synthesis pathway mutant *cha-1* was used as a control as it is reported to show resistance to aldicarb (Rand and Russell, 1984). These animals when exposed to aldicarb and EtOH did not show the defect in movement seen in WT *C. elegans* (Figs. 6a and b). As a control *cha-1* mutants were also analyzed with just EtOH (no aldicarb) and these animals also behaved in a manner similar to that of WT control animals (Fig. S5). These experiments indicate that the *dop-2* EIS behavior is an outcome of increased cholinergic signaling at the NMJ. ACh signals mainly through the nicotinic ACR-16 receptors present on the postsynaptic body-wall muscle membrane. In *acr-16* mutant there is an 85% decrease in response to ACh when compared to WT animals (Touroutine et al., 2005). To understand the role of the acetylcholine receptors in the EIS behavior, we tested a previously used ACR-16::GFP line for the EIS behavior (Babu et al., 2011). We found that overexpression of ACR-16 caused the animals to show the EIS behavior while mutants in *acr-16* behaved like WT animals after EtOH treatment (Figs. 6c and d). However, the phenotype seen on ACR-16 overexpression was not as robust as that seen with *dop-2* mutant animals. Together these data indicate that elevated muscle excitation through increased acetylcholine is playing a significant role in the EIS behavior of the animal.

**Fig. 6:**
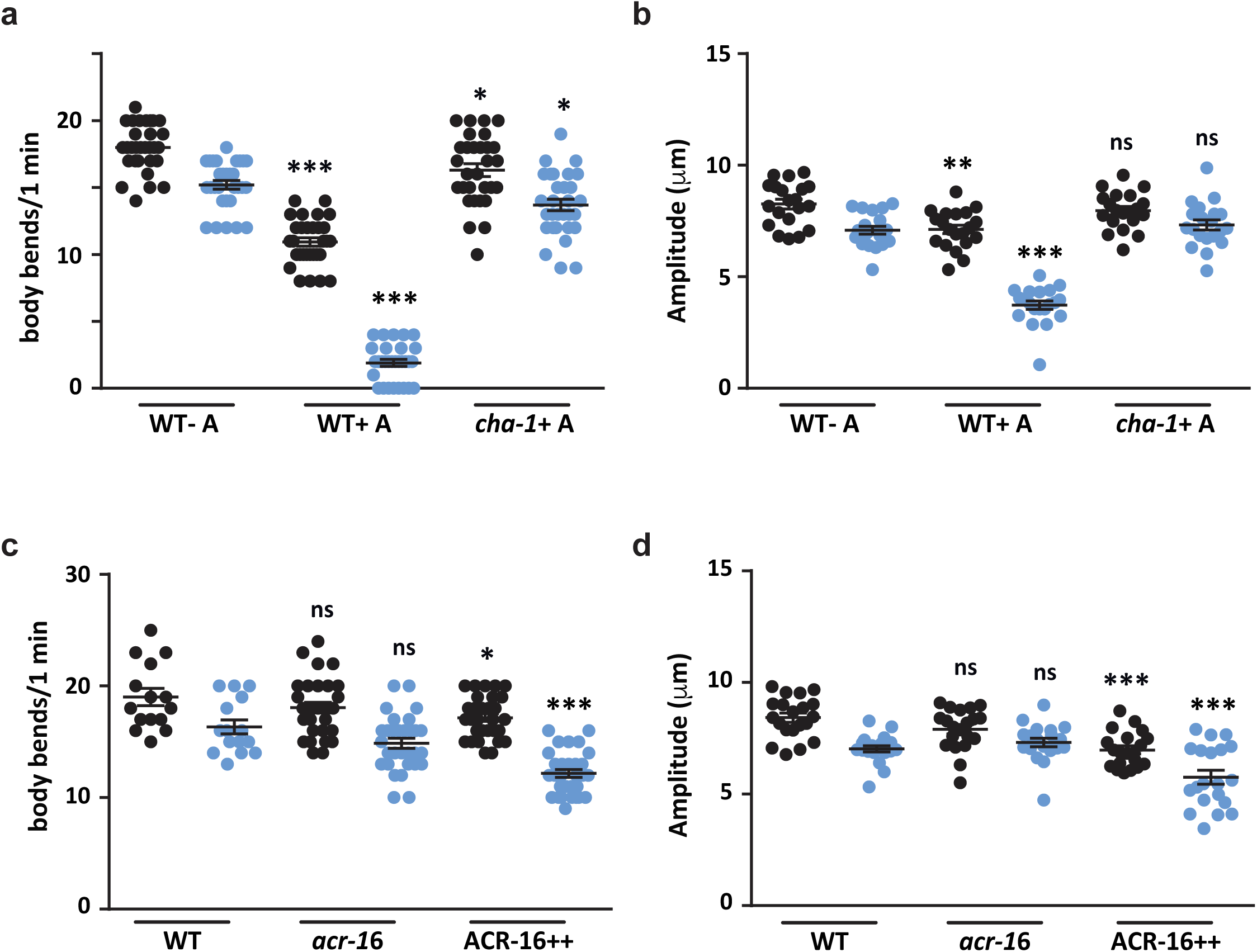
Increased acetylcholine signaling causes increased sensitivity to ethanol in WT animals. (a) Graph of number of body-bends for WT and *cha-1* mutants on EtOH plates after treatment with aldicarb, (F = 254, DF = 5). (b) Graph indicates amplitude of body-bends for WT and *cha-1* mutants on EtOH plates after treatment with aldicarb, (F = 68.6, DF = 5). (c) Quantitation of number of body-bends from WT, *acr-16* and the ACR-16++ line on EtOH, (F = 28.2, DF = 5). (d) Quantitation of amplitude of body-bends from WT, *acr-16* and the ACR-16++ line on EtOH, (F = 18.4, DF = 5). Error bars represent ±S.E.M. and p-values were calculated using one-way ANOVA and Turkey-Kramer multiple comparison test; “*” indicates p<0.05, “**” indicates p<0.01, “***” indicates p<0.001 and “ns” indicates not significant in all panels.

## Discussion

DOP-2 belongs to a family of D2-like inhibitory receptors. It is thought to negatively regulate the release of dopamine by feedback inhibition of dopamine release from presynaptic neurons (Reviewed in (De Mei et al., 2009; Ford, 2014; Mercuri et al., 1997)). In *C. elegans* DOP-2 is only present on the DA neurons making it a very good candidate for autoregulation of DA. However, its deletion does not show defects in DA dependent behaviours as reported for other DA receptors such as DOP-1 and DOP-3 (Allen et al., 2011; Chase et al., 2004; Sawin et al., 2000). It is likely that behaviors associated with deletion of neuromodulatory molecules are not easily observable in native conditions since they are required to modulate multiple behaviors and not one specific behavior (Ford, 2014). EtOH has been shown to increase dopamine release from the mammalian ventral tegmental area and increased dopamine levels were found in the nucleus acccumens (Imperato and Di Chiara, 1986; Weiss et al., 1996; Yim and Gonzales, 2000). Since both loss of D2-like receptors and ethanol (EtOH) tend to increase DA levels, we went on to test *dop-2* deletion mutants for movement defect/s in the presence of EtOH and found a robust behavior involving decrease in body-bends and the flattening of the body-bends which we have termed Ethanol Induced Sedative (EIS) behavior. This behavior was more pronounced in the posterior region of the animal. Although previous studies have reported that *C. elegans* show tolerance towards acute EtOH exposure and after recovery they exhibit various forms of disinhibitions in their behaviors (Mitchell et al., 2010; Topper et al., 2014), our studies describe for the first time the role of chronic EtOH treatment for extended periods of time (400 mM EtOH for upto 24 hrs). Our data also shows that the EIS behavior in *dop-2* mutant animals is due to increased dopamine release.

The synaptic levels of dopamine are maintained by the activities of dopamine transporter (DAT-1), that recycles the dopamine back to the cell in conjunction with the DA autoreceptor (DOP-2) (Benoit-Marand et al., 2000; Schmitz et al., 2002). In mammals the activity of the DA transporter can be controlled by D2-autoreceptors by regulating their surface expression (Benoit-Marand et al., 2011; Cass and Gerhardt, 1994; Dickinson et al., 1999; Mayfield and Zahniser, 2001; Schmitz et al., 2002; Wu et al., 2002). This occurs at least partially via an increase in DAT cell surface expression after D2-receptor activation (Mayfield and Zahniser, 2001). However, D2-antagonists or D2 deletion could not alter DAT dependent DA uptake (Anzalone et al., 2012; Beckstead and Williams, 2007; Bello et al., 2011; Benoit-Marand et al., 2011; Benoit-Marand et al., 2001; Kennedy et al., 1992). Our FRAP experiments conducted in the presence of increased EtOH showed increased release of DA and in the absence of *dop-2*. Further the membrane expression of DAT-1 was decreased in EtOH treated *dop-2* mutant animals when compared to control animals. Hence we show that DAT-1 surface transport is dependent upon DOP-2. However, since loss of *dat-1* does not show the EIS phenotype, it is likely that multiple factors including increased dopamine release contribute to the EIS behavior and it is not dependent on just loss of DAT-1 membrane expression.

All our data points towards the fact that the posterior region of the animal is more affected then the anterior region during EIS behavior in *dop-2* mutants. Neuronal ablation experiments demonstrate that the posterior DA neuron, PDE is responsible for the *dop-2* EIS phenotype. The PDE neuron forms multiple unidirectional synapses with the DVA interneuron. Our experiments implicate PDE function through the DVA neuron to allow for changes in downstream motor circuitry. These findings are in line with work that has previously reported that the PDE neuron makes direct synaptic contacts with the DVA neuron (Bhattacharya et al., 2014). DVA is known to modulate locomotion both positively and negatively by providing a unique mechanism whereby a single neuron can fine-tune motor activity (Li et al., 2006). The DVA interneuron has connections with both motor neurons and interneurons and relays information for normal locomotion (Bhattacharya et al., 2014; Gray et al., 2005). One mechanism of maintaining normal locomotion especially in conditions of stress like EtOH exposure could involve dopamine release from PDE regulating the movement of *C. elegans* through DVA. Our work further implicates the role of the DOP-1 receptor in DVA and the requirement of this pathway to maintain the EIS behavior seen in *dop-2* mutants. The DVA neuron expresses the DA receptor DOP-1, a D1-like excitatory receptor (Bhattacharya et al., 2014). Further, in *Drosophila* and mammals it has been demonstrated that D1-like DA receptors promote EtOH-induced disinhibition (Abrahao et al., 2011; Kong et al., 2010).

A prior study has shown that movement induces NLP-12 release from DVA neurons and enhances ACh release at NMJs (Hu et al., 2011). Further, multiple studies have demonstrated that neuropeptides modulate neuronal activity and synaptic transmission (Bhardwaj et al., 2018; Edwards et al., 2009; Hu et al., 2011; Jacob and Kaplan, 2003; Kass et al., 2001; Sieburth et al., 2007; Speese et al., 2007; Sumakovic et al., 2009). In this study we show that overexpressing just the NLP-12 neuropeptide in the DVA interneuron is sufficient to mimic the EIS phenotype seen in *dop-2* mutants, again implicating neuropeptides in synaptic functions.

Previous work that has shown that in *Drosophila melanogaster*, mutants in a gene called *arouser* cause the animal to show increased sedation in the presence of EtOH as well as show increased boutons at the NMJ, indicating a link between increased neuromuscular signaling and greater susceptibility to alcohol (Eddison et al., 2011). Our data demonstrates that the EIS behavior seen in *dop-2* mutants could occur because of changes in neurotransmission at the *C. elegans* NMJ. We also show that increased neurotransmission brought about by the drug aldicarb or overexpressing acetycholine receptors at the body-wall muscle both cause EIS behavior in *C. elegans*.

Overall our data implicates *dop-2* in the ethanol induced sedative phenotype by showing that 1. Mutants in *dop-2* show increased dopamine release in the PDE neuron upon chronic ethanol treatment. 2. The increased dopamine released from the PDE sensory neuron causes increased release of the neuropeptide, NLP-12 from the DVA interneuron. 3. Increased NLP-12 release could cause increased cholinergic neuronal function and result in the EIS phenotype observed. This circuit is illustrated in Fig. 7.

**Fig. 7:**
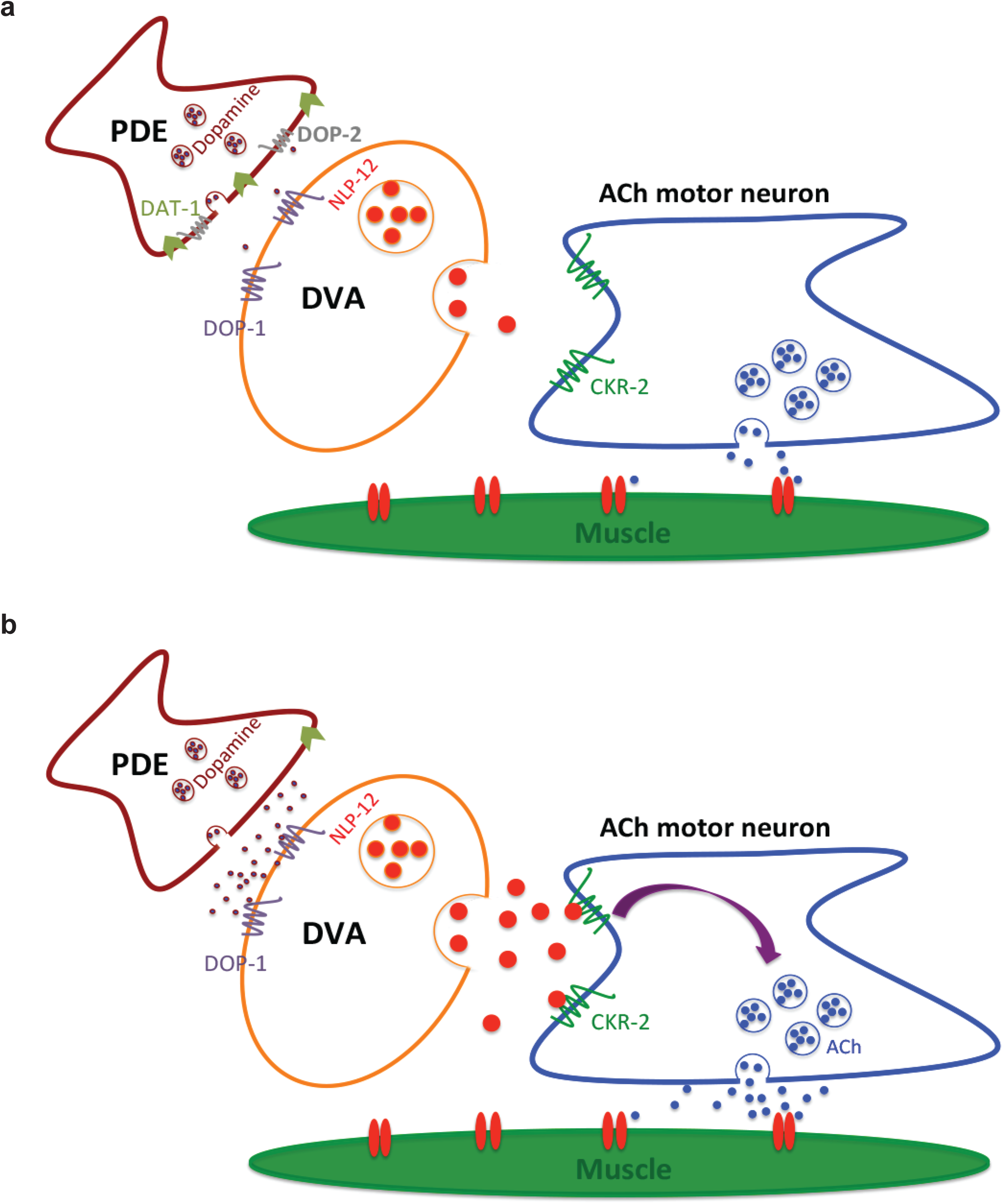
Proposed model for DOP-2 functioning in the presence of EtOH. (a) DOP-2 DA autoreceptors functions in the posterior DA neurons (PDE) to regulate DA levels, in the presence of EtOH. (b) Loss of the *dop-2* autoreceptor leads to unregulated release of DA in the presence of EtOH. The increased levels of DA activate the function of DOP-1 receptors present on the DVA neuron causing increased DVA activation. This in turn causes increased release of the neuropeptide NLP-12, which in turn could cause increased cholinergic signaling at the NMJ. This model is based on this work and prior studies (Bhattacharya et al., 2014; Hu et al., 2011).

Taken in its entirety, our work along with previous work paves a path for using *C. elegans* as a model system to study the molecular players involved in alcohol dependent locomotory functions (Davies et al., 2003; Mitchell et al., 2007).

## Methods

### Strains

Animals were maintained according to standard protocols (Brenner, 1974). N2 Bristol was used as the wild type (WT) strain. The mutant strains used in this study were; *dop-2(vs105), dop-3(vs106), dop-1(vs100), cat-2(n4547), cha-1(p1152), acr-16(ok789), slo-1(eg142), dat-1(ok157)* and *nlp-12(ok335)*.

### Constructs and transgenes

All constructs were generated using *pPD95.75* as the backbone with standard restriction digestion cloning procedures (Russell, 2001). Transgenic lines were generated by microinjection of the desired plasmid as previously described (Mello and Fire, 1995). The *Pdop-2*::DOP-2::mCherry construct was obtained from Rene Garcia Lab (Correa et al., 2012). The *nlp-12* promoter used in pBAB911 was cloned by amplifying a 355 bp upstream region of the *nlp-12* gene from genomic DNA using AACTGCAGGGCCGAGACGAATCCGGAGG (AS1) and CGGGATCCGCATTTTGTCGGAGGCAATT (AS4) primers and cloned into the *pPD95.75* vector using Pst I and Bam HI sites. The pBAB913 construct was generated by cloning P*nlp-12* into a previously made P*exp-1*::*sl2*::wrmScarlet vector using Pst I and Xma I sites that removed P*exp-1*. The pBAB912 construct contains a 1.7 kb genomic region of *nlp-12* amplified from genomic DNA using AACTGCAGGGCCGAGACGAATCCGGAGG (AS1) and CGGGATCCGAAAATGTGTCGCTTCGAGAC (AS3) primers. The PCR amplified fragment was cloned into the *pPD95.75* vector using Pst I and Bam HI sites, for generating the *nlp-12* overexpression lines. For *dop-1* rescue experiment *dop-1* cDNA (1.2kb) was cloned under the *nlp-12* promoter in pBAB913 using Xma I and Kpn I sites. A complete list of strains, primers and plasmids used in this study are available in the supplementary tables S1, S2 and S3 respectively.

### Ethanol induced behavioral assay

*C. elegans* were synchronized by bleaching and grown on nematode growth media (NGM) plates. They were maintained in well-fed conditions till the assay. The Ethanol (EtOH) plates were prepared using unseeded plates i.e. without food plates that were dried for 3 hours (h) under airflow in the biosafety cabinet. This was followed by spreading EtOH (400 mM) on the plates. The plates were then sealed with parafilm and were allowed to equilibrate for 2 h at 20°C. Adult animals were initially transferred to unseeded plates for 15 to 20 seconds (sec) and then moved to the EtOH plates. 10 animals were used for each set and the experiment was done in triplicate for every genotype tested. All animals initially showed coiling behavior and paralyzed within 10 minutes (min) of EtOH exposure as has been previously reported (Davies et al., 2003). Recovery of *C. elegans* started in all the strains except *dop-2* in approximately 30 min. All the strains used for assays including WT showed Ethanol Induced Sedentary (EIS) - like behavior at around 30 min when movement started. However, most strains regained their normal locomotory behavior after more then 60 min. The plates were incubated overnight (16-20 h) to visualize the tracks. The quantitative analysis of behavior was done at 120 min for all the strains by recording the videos10 frames/sec for 1 min on the AxioCam MRm (Carl Zeiss) using the micromanager software. The anterior and posterior body-bends and amplitude of body-bends were quantified separately for each animal using the imageJ software (Schindelin et al., 2012), n=30 animals were observed for each genotype for body-bends and n=20 for the amplitude of body-bends. Videos obtained from the same animal were used to quantify both body-bends and amplitude of locomotion. The results were plotted as graphs using GraphPad Prism v6 and statistics were evaluated using one-way ANOVA.

### Exogenous Dopamine Assay

For exogenous dopamine application, 1 M freshly prepared dopamine was used as previously described (Baidya et al., 2014). The final concentration of dopamine used was 40 mM. Dopamine was spread on EtOH plates for EtOH + dopamine experiments and on unseeded dry plates for only dopamine exposure experiments. The plates were protected from light and used within 10 min of preparation for each assay. Animals were transferred from unseeded plates to the assay plate and their behavior was analyzed by making 1 min videos at 10 frames/sec after 2 h of dopamine/EtOH exposure.

### Aldicarb treatment prior to the EtOH assay

This assay was modified from the aldicarb assay to study the behavior of *C. elegans* on EtOH exposure after aldicarb treatment. The aldicarb assay was performed as described previously (Mahoney et al., 2006; Sieburth et al., 2005; Vashlishan et al., 2008). Briefly, plates with 1 mM aldicarb (Sigma-Aldrich 33386) were prepared 1 day prior to the assay and dried. *C. elegans* were transferred to aldicarb plates for 60 min and then moved to EtOH plates after which they were analyzed as previously described in the EtOH induced behavioral assay section.

### Microscopy

The DAT-1::GFP imaging experiment was performed on the Leica SP8 confocal microscope using the Argon laser at 10% gain. Young adult animals were immobilized using 30 mg/ml BDM (2,3-ButaneDione monoxime) on 2% agarose. All the image quantitation was done taking whole cell body expression of GFP using FIJI. Experiments were performed both with and without EtOH exposure.

### Neuronal ablation

The ablation of PDE and DVA neurons were done using Bruker Corporations ULTIMA two photon IR laser system. In which, one laser was used for imaging (920nm for GFP, 1040nm for mScarlet) and another laser for ablating the neurons (720 nm, irradiation duration-20ms, pulse width 80fs, power∼23mW) as described in (Basu et al., 2017). During ablation, L2 staged worms were immobilized on 5% agarose pads using 10mM levamisole hydrochloride (Sigma-Aldrich 10380000) or 0.1 μm-diameter polystyrene beads (00876-15; Polystyrene suspension). These worms were then recovered using mouth pipette onto the newly seeded NGM plates and were allowed to grow uptill young adult stage. The worms were then evaluated in the EtOH induced behavioral assay.

### FRAP experiments

The increased extracellular release kinematics of dopamine in the presence of EtOH based on DA vesicle fusion, was analyzed and observed using the dopaminergic promoter (P*asic-1*) tagged with pH sensitive synaptobrevin-super ecliptic pHluorin reporter fusion construct (SNB-1::SEpHluorin) (Hardaway et al., 2015; Samuel et al., 2003; Voglis and Tavernarakis, 2008). Young adult animals were mounted on 2% agarose pads and paralyzed using 0.05% levamisole hydrochloride (Sigma-Aldrich 10380000). FRAP experiments were performed on the Leica SP8 inverted confocal microscope. PDE synapses were identified by Synaptobrevin::SEpHluorin fluorescence. Bleaching was done using the 488 nm argon laser, 80% bleach power for 5-10 sec to an intensity of 10-15% of original fluorescence value. Fluorescence was monitored every 10 sec for 2 min and analyzed using FIJI software. The percentage recovery was calculated at each time point and 20 synapses were analyzed per genotype. The data was plotted using GraphPad Prism v6 and analyzed using non-linear regression plotting and one phase association exponential equation was used to analyze this data.

### Statistical analysis

All statistical analyses were performed by using GraphPad Prism Version 6.0. The error bars represent SEM. Statistical comparisons were done using one-way ANOVA with Turkey-Kramer multiple comparison test. The level of significance was set as “*” indicates p < 0.05, “**” indicates p < 0.01 and “***” indicates p < 0.001.

## Acknowledgments

The authors are especially grateful to Yogesh Dahiya for help with the FRAP experiment. We thank Randy Blakely, Rene Garcia and Ron Evans for reagents. A number of strains were provided by CGC, which is funded by NIH Office of Research Infrastructure Programs (P40 OD010440). The authors thank Ankit Negi for routine help and the IISER Mohali Confocal facility for use of the confocal microscope.

AS thanks Council of Scientific and Industrial Research (CSIR)-University Grants Commision (UGC) for a graduate fellowship. PP acknowledges support from a Department of Science and Technology (DST)-Woman of Science (WOS-A) grant as well as past funding from Department of Biotechnology (DBT) Bio-CARe, Indian Institute of Science Education and Research (IISER) Mohali and an IA grant awarded to KB. KB was an Intermediate Fellow of the India Alliance (IA) and thanks the Alliance for funding support. KB also thanks DST-Science and Engineering Research Board (SERB) for funding support.

## Funding

This work was supported by the Wellcome Trust/ DBT India Alliance Fellowship [grant number IA/I/12/1/500516] awarded to KB and partially supported by a DST– SERB grant [SERB/F/7047] to KB. PP is supported by a DST WOS-A grant [SR/WOS-A/LS-285/2018] and was earlier supported by a DBT Bio-CARe grant [BioCARe/01/10167]. AG-R lab is supported by the NBRC core fund from the Department of Biotechnology and Wellcome Trust-DBT India Alliance (Grant # IA/I/13/1/500874).

## Author Contributions

PP and AS designed, performed, analyzed all the experiments and wrote the manuscript. PP and AS are co-first authors listed alphabetically. HK and AG-R helped with performing the ablation experiments and editing the manuscript. KB supervised the experiments, helped with experimental design and data interpretation and edited the manuscript.

The authors declare no conflict of interest.

## Supplementary tables

**Fig. S1:**
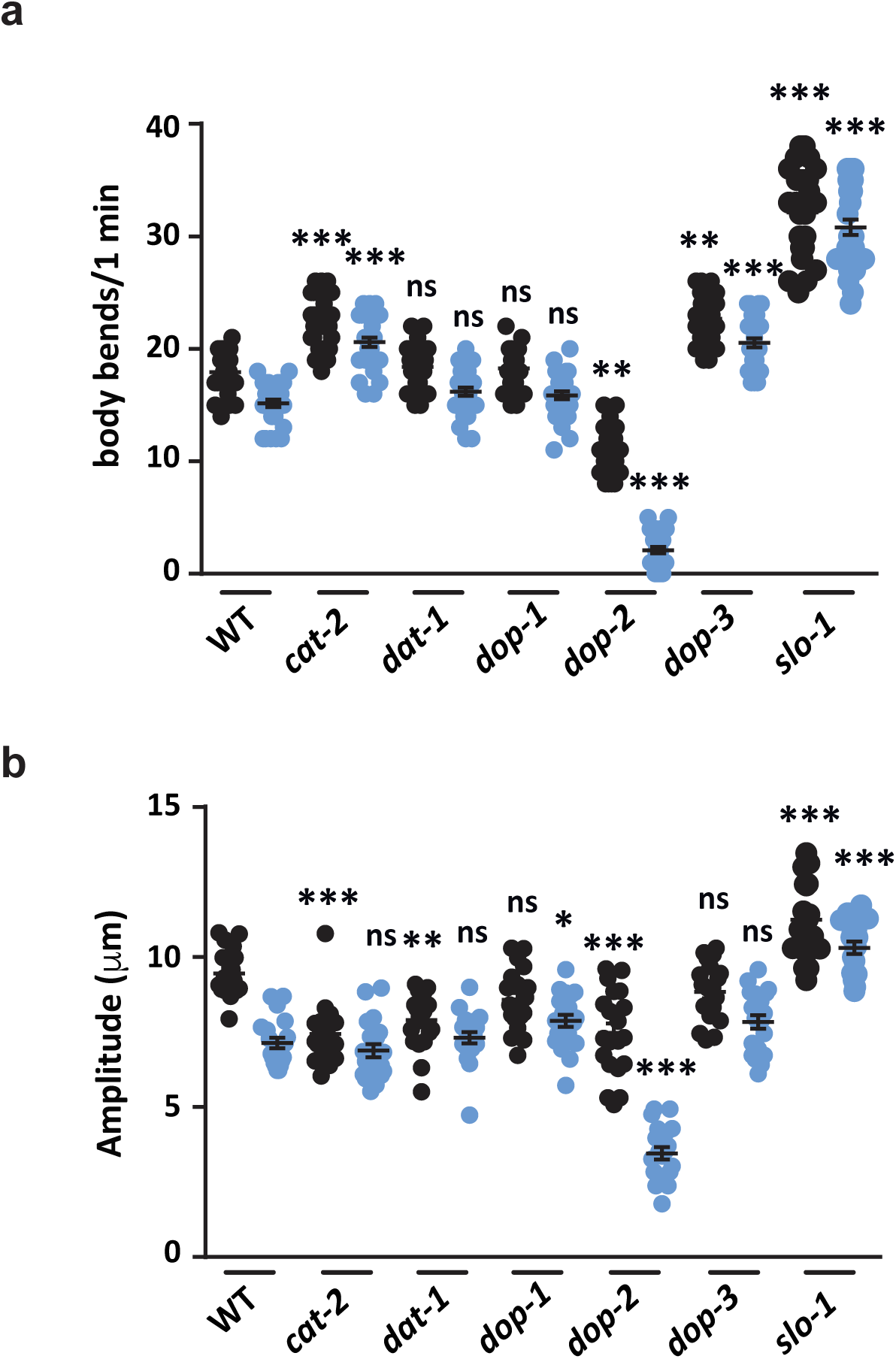
Ethanol dependent phenotype of dopaminergic pathway mutants. (a) Graph of number of body-bends (posterior body-bends are shown in blue) in WT, *cat-2*, *dat-1*, *dop-1*, *dop-2*, *dop-3* and *slo-1* mutants upon EtOH treatment. Experiments were performed in triplicates with 10 worms per assay in all the figures. For this graph F = 66.8 and DF = 13. (b) Graph of amplitude of body-bends (posterior body-bends are shown in blue) for WT, *cat-2*, *dat-1*, *dop-1*, *dop-2*, *dop-3* and *slo-1* mutants upon EtOH treatment. Ten animals were used for each experiment to quantitate the amplitude of body-bends and the experiment was performed in duplicate for all figures. The same videos of moving animals were used to quantitate both number and amplitude of body-bends. For this graph F = 306 and DF = 13. In both graphs mutants in the EtOH sensing channel protein, *slo-1* were used as positive control. Error bars represent ±S.E.M. and p-values were calculated using one-way ANOVA and Turkey-Kramer multiple comparison test; “*” indicates p<0.05, “**” indicates p<0.01, “***” indicates p<0.001 and “ns” indicates not significant in all panels.

**Fig. S2:**
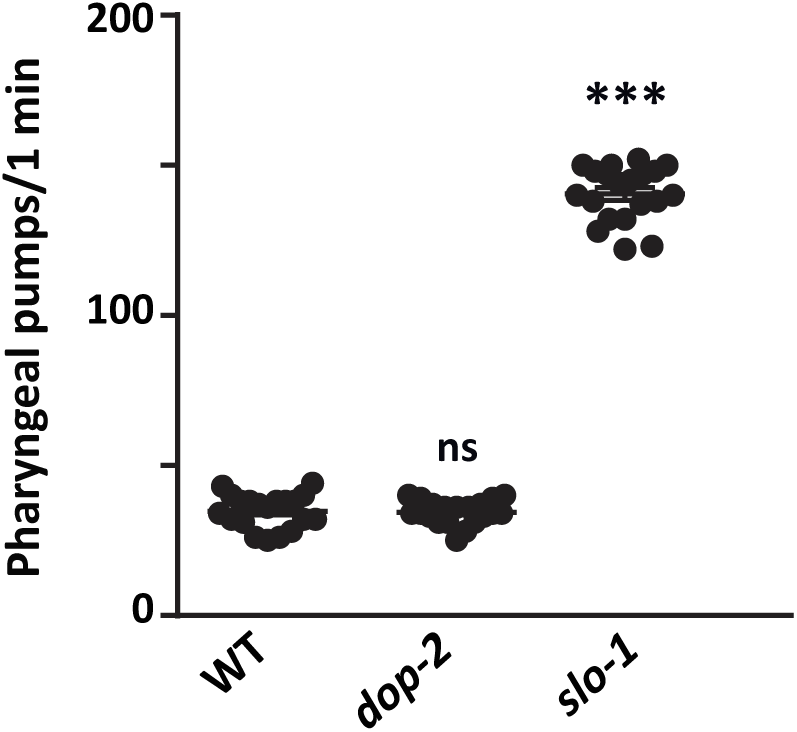
Mutants in *dop-2* do not show defects in pharyngeal pumping. The graph shows the quantitation of the number of pharyngeal pumps in 1 min from WT, *dop-2* and *slo-1* mutants in the presence of EtOH. The experiment was performed with 10 animals per experiment in duplicate. Error bars represent ±S.E.M. and p-values were calculated using one-way ANOVA and Turkey-Kramer multiple comparison test; “***” indicates p<0.001 and “ns” indicates not significant.

**Fig. S3:**
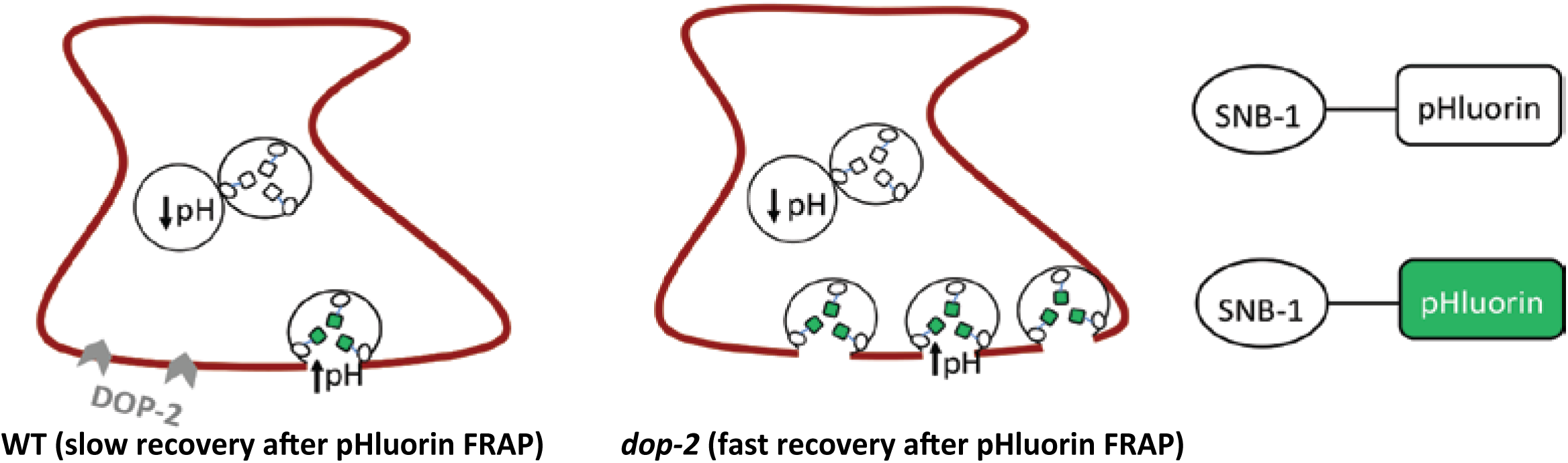
Diagrammatic representation of the FRAP experiment. A pH sensitive GFP Fluorophore, pHluorin, was tagged to the synaptic vesicle protein SNB-1 and expressed in DA neurons. This line was constructed and previously used (Hardaway et al., 2015).

**Fig. S4:**
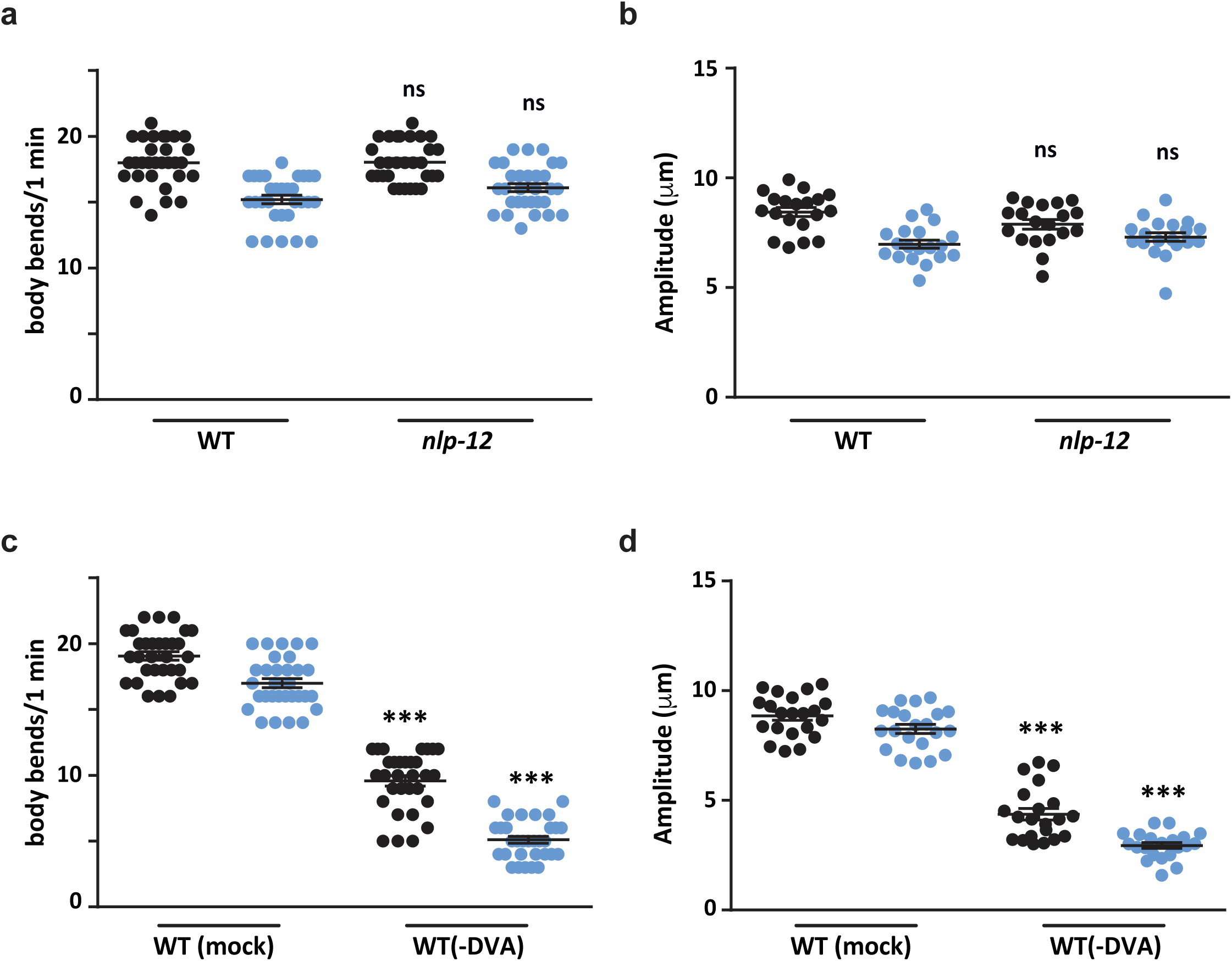
DVA ablation affects locomotion in WT animals. (a) Quantitation of the number of body-bends from WT and *nlp-12* mutant animals upon EtOH treatment, (F = 12.2, DF = 3). (b) Quantitation of the amplitude of body-bends from WT and *nlp-12* mutants upon EtOH treatment, (F = 10.1, DF = 3). (c) Graph represents number of body-bends in mock and DVA ablated WT animals, 10 animals were used for the mock treatment and 11 for the DVA ablation. The experiment was performed in triplicate, (F = 368, DF = 3). (d) Graph shows the amplitude of body-bends in mock and DVA ablated WT animals, 10 animals were used for the mock treatment and 11 for the DVA ablation. The experiment was performed in duplicate (F = 197, DF = 3). Error bars represent ±S.E.M. and p-values were calculated using one-way ANOVA and Turkey-Kramer multiple comparison test; “***” indicates p<0.001 and “ns” indicates not significant.

**Fig. S5:**
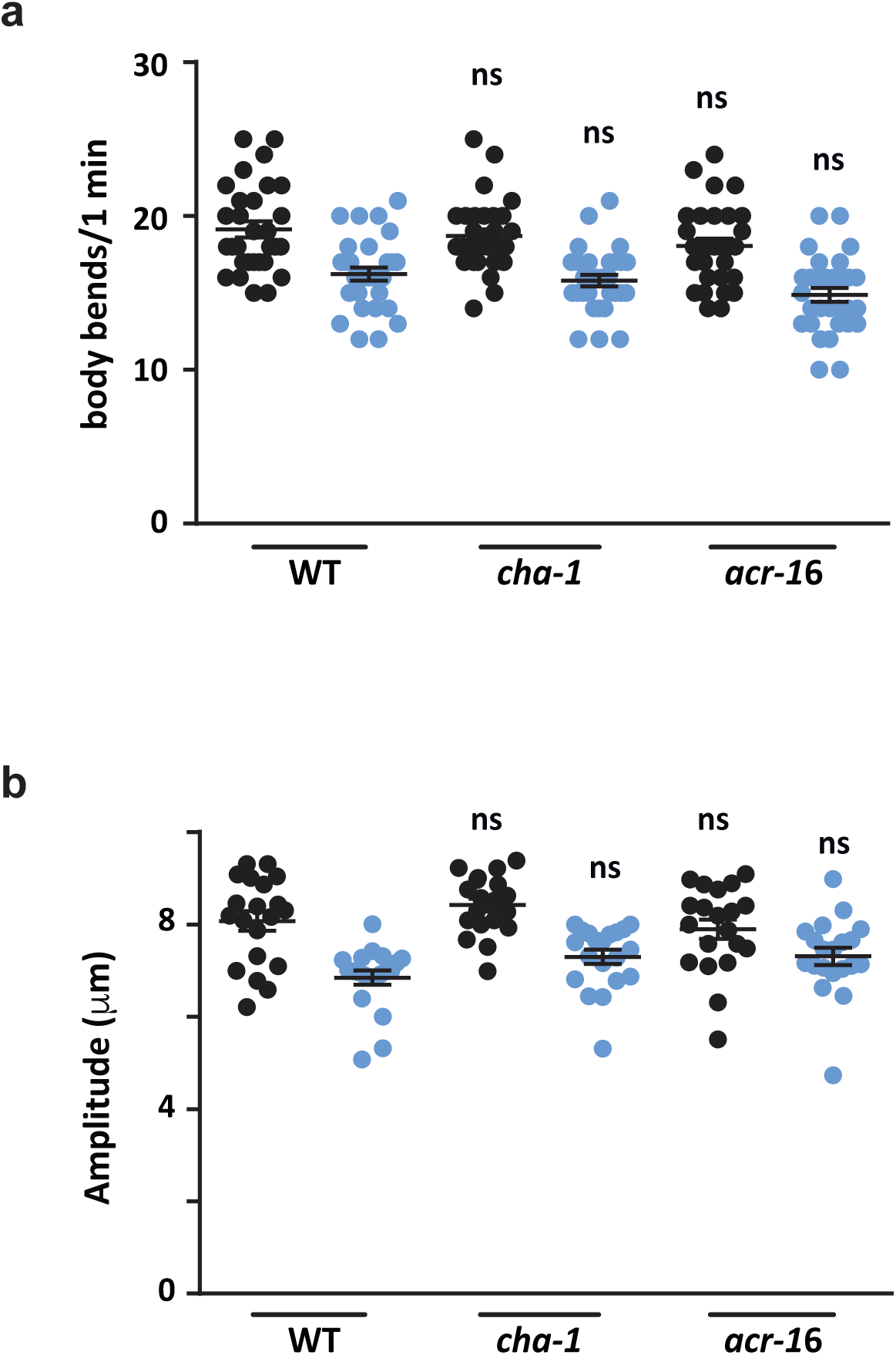
Mutants in the cholinergic pathway do not show a phenotype in the presence of EtOH. (a) Graph indicates the number of body-bends quantitated from WT, *cha-1* and *acr-16* mutants upon EtOH treatment, (F = 15, DF = 5). (b) Graph shows amplitude of body-bends quantitated from WT, *cha-1* and *acr-16* mutants upon EtOH treatment, (F = 10.8, DF = 5). Error bars represent ±S.E.M. and p-values were calculated using one-way ANOVA and Turkey-Kramer multiple comparison test; “ns” indicates not significant in both panels.

**Table S1:** List of strains used in this study.

| Strain | Genotype | Comments |
| --- | --- | --- |
| **LX702** | *dop-2(vs105)V* | CGC strain |
| **LX703** | *dop-3(vs106)X* | CGC strain |
| **LX645** | *dop-1(vs100)X* | CGC strain |
| **MT15620** | *cat-2(n4547)II* | CGC strain |
| **PR1152** | *cha-1(p1152)IV* | CGC strain |
| **RB918** | *acr-16(ok789)V* | CGC strain |
| **BZ142** | *slo-1(eg142)V* | CGC strain |
| **RM2702** | *dat-1(ok157)III* | CGC strain |
| **RB607** | *nlp-12(ok335)* | CGC strain |
| **nuls299** | *Pmyo-3::ACR-16::GFP* | Josh Kaplan Lab |
| **IR724** | *asic-1::SNB-1::SEpHlourin* | Blakely Lab |
| **BY834** | *Pdat-1::DAT-1::GFP* | Blakely Lab |
| **BZ555** | *egls1(Pdat-1::gfp)* | CGC strain |
| **BAB900** | *dop-1(vs100); dop-2(vs105)* | This study |
| **BAB901** | *cat-2(n4547); dop-2(vs105)* | This study |
| **BAB902** | *egls1(Pdat-1::gfp); dop-2(vs105)* | This study |
| **BAB903** | *Pdat-1::DAT-1::GFP; dop-2(vs105)* | This study |
| **BAB904** | *asic-1::SNB-1::SEpHlourin; dop-2(vs105)* | This study |
| **BAB905** | *Pdop-2::DOP-2::CFP; dop-2 (IndEx905)* | This study |
| **BAB906** | *Pnlp-12::NLP-12 (IndEx906)* | This study |
| **BAB907** | *Pnlp-12::DOP-1::wrm Scarlet; dop-1(vs100);* | This study |
|  | *dop-2(vs105)* (IndEx907) |  |
| **BAB908** | *Pnlp-12::DOP-1::wrm* Scarlet; *dop-1(vs100)*<br>(IndEx908) | This study |

**Table S2:**
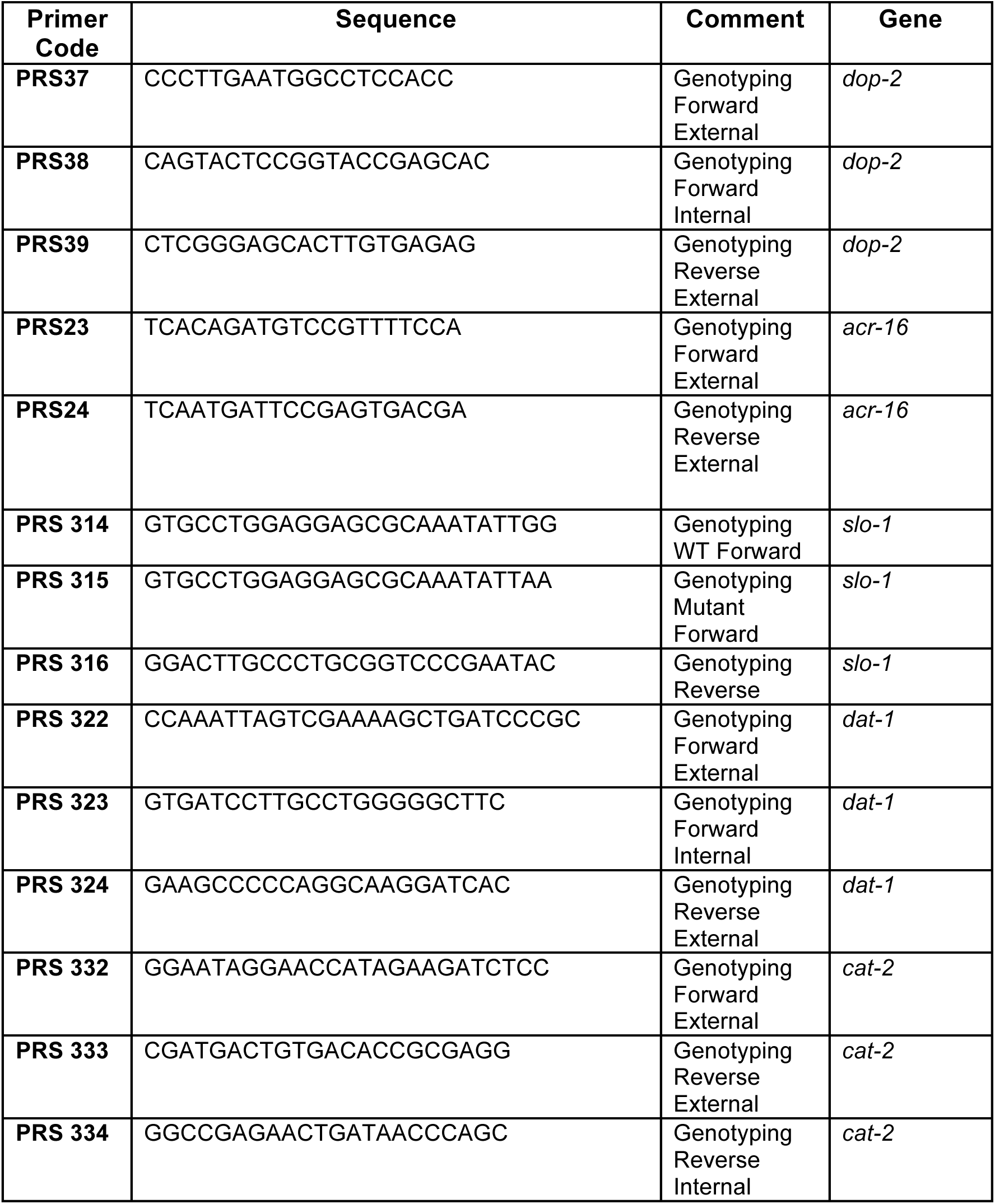
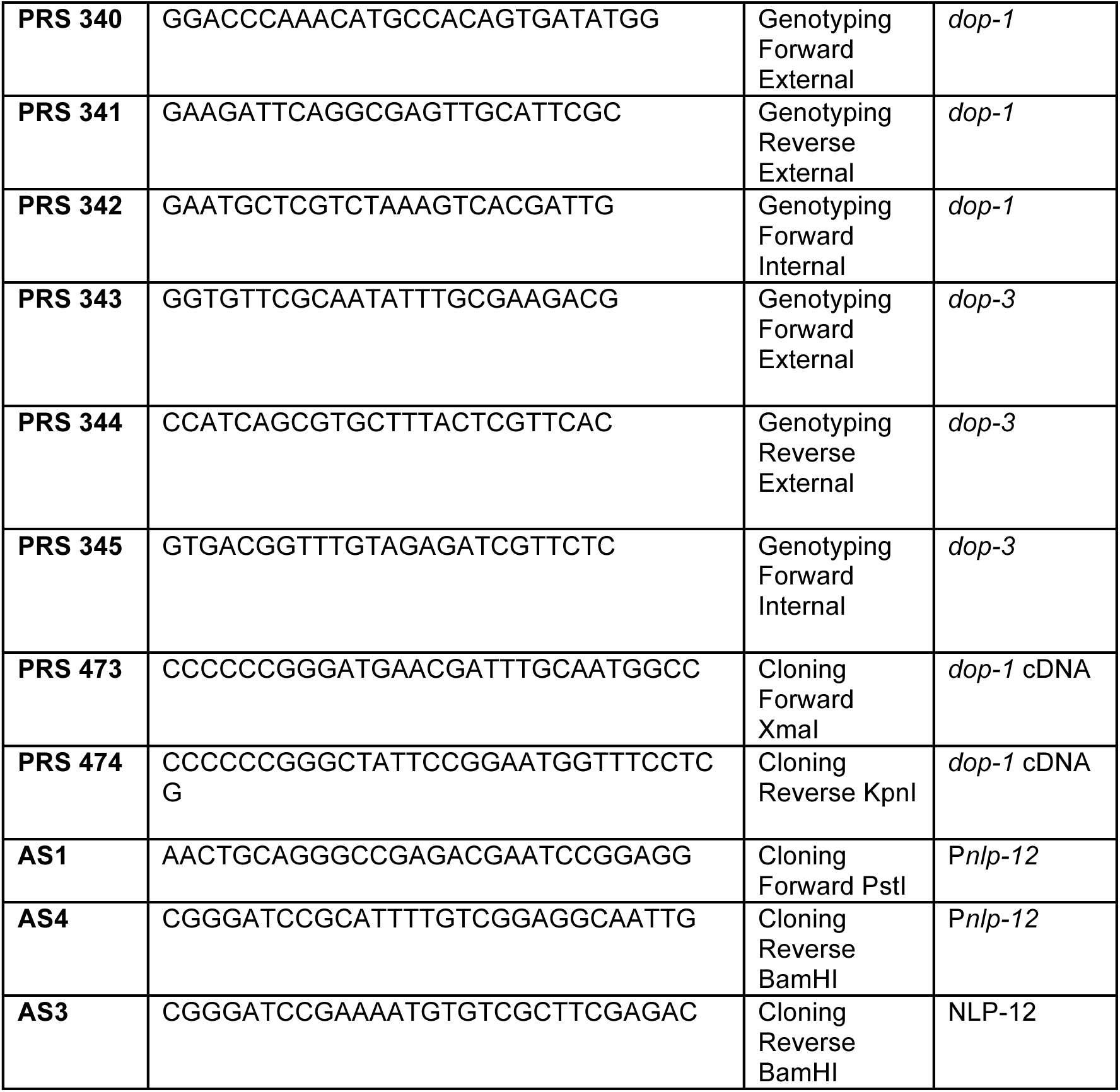
List of primers used in this study.

**Table S3:**
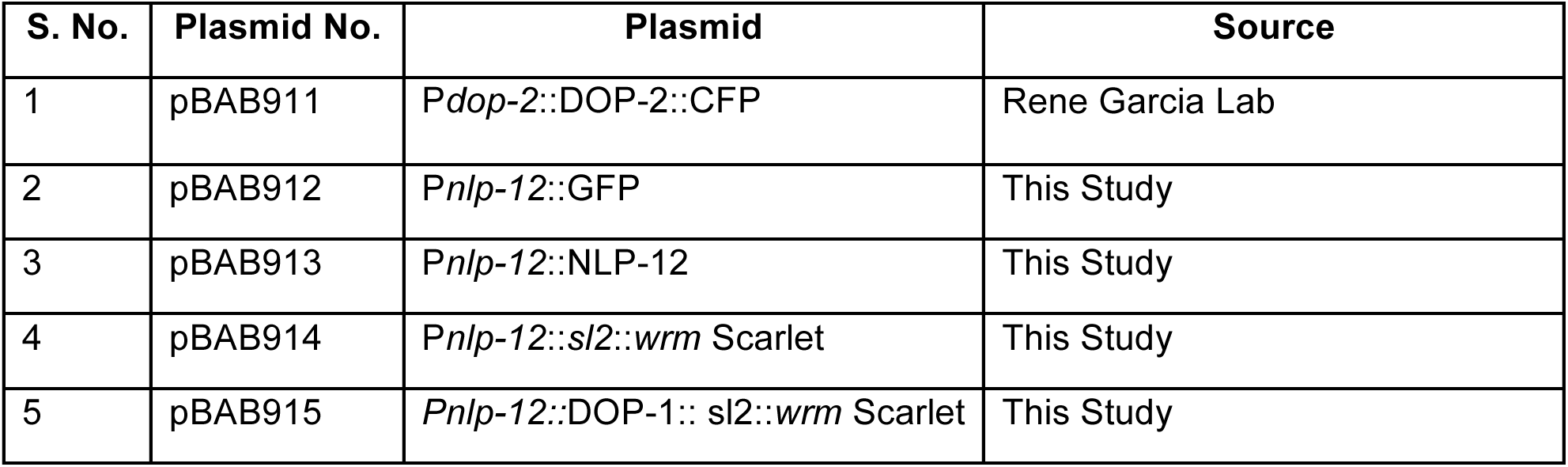
List of plasmids used in this study.

